# SorCS1 promotes synaptic and cognitive resilience despite amyloid pathology in Alzheimer’s disease model mice

**DOI:** 10.64898/2026.08.24.746580

**Authors:** Nayoung Yi, Alfred Kihoon Lee, Farin B. Bourojeni, Manni Wang, Mai Inagaki, Hideto Takahashi

## Abstract

Alzheimer’s disease (AD) lacks effective therapies despite extensive efforts targeting amyloid-β (Aβ) and its precursor processing. Synapse loss is the strongest correlate of cognitive decline, driven partly by Aβ oligomers (AβOs), which bind multiple synaptic membrane proteins including the synaptic organizer neurexin and disrupt synaptic integrity and function. The protein-sorting receptor SorCS1 blocks interactions between AβOs and β-isoforms of neurexins (β-Nrxns), but its therapeutic relevance *in vivo* remains unclear. Using 5xFAD mice, which overproduce AβOs, combined with forebrain-specific neuronal SorCS1 overexpression, we show that SorCS1 preserves working memory, synaptic integrity, and basal excitatory transmission without altering amyloid deposition, in part by restoring synaptic β-Nrxn expression. SorCS1 also reduces tau hyperphosphorylation in 5xFAD synaptosomes and binds the tau kinase GSK3β. These results identify SorCS1 as an AD resilience-promoting factor that maintains synaptic connectivity and attenuates tau pathology, revealing a therapeutic strategy that operates independently of amyloid reduction.

## Introduction

Alzheimer’s disease (AD) is the most common form of dementia and a major public health challenge in aging societies (*1*). The neuropathological hallmarks of AD are well established: senile plaques, which are extracellular deposits primarily composed of toxic amyloid-β (Aβ) peptides, and neurofibrillary tangles, which are intracellular aggregates of hyperphosphorylated tau proteins (*2, 3*). Despite decades of intensive research, therapeutic progress has been limited (*4, 5*). Numerous clinical trials have targeted Aβ peptides, their aggregates, or the processing pathways of the amyloid precursor protein (APP), yet none have yielded effective or durable clinical outcomes (*6*). This lack of success highlights the urgent need to identify novel molecular mechanisms underlying AD pathology that could serve as the basis for innovative therapeutic strategies.

Synaptic pathology is a central feature of AD and is associated more strongly with cognitive decline than plaque or tangle burden (*7, 8*). Postmortem and biopsy studies have shown that synapse loss is tightly correlated with impaired cognition (*9*). Experimental evidence indicates that soluble Aβ oligomers (AβOs) disrupt synaptic function and drive synapse loss *in vitro* and in AD model mice (*10*). On the other hand, biomarker studies have suggested that Aβ accumulation begins years before synaptic loss, implying that synapses can tolerate Aβ under certain conditions (*11*). The concept of “cognitive resilience” supports this view: some individuals maintain normal cognition despite advanced AD pathology, and their synaptic protein expression resembles that of healthy controls (*12–15*). These findings suggest that synapses may exist in states of vulnerability or tolerance to Aβ. The molecular basis of this distinction remains unclear, and its elucidation is critical for understanding disease progression and developing strategies to preserve synapse structure and function in AD.

Synaptic organizers are strong candidates for mediating Aβ vulnerability and tolerance (*16*). These cell adhesion molecules regulate synapse formation, maturation, and plasticity, and are essential for cognitive function. Many synaptic organizing complexes have been identified (*17, 18*), including neurexin (Nrxns)–neuroligin (Nlgn), Nrxn-LRRTM1/2, PTPσ-TrkC, and PTPδ-Slitrk. Our group has shown that Nrxns interact directly with AβOs leading to reduced Nrxn surface expression and impaired excitatory synaptic organization (*19*). Human studies have reported reduced expression of NRXN3 and NLGN1 in AD brains (*20–22*), whereas NRXN1 and NLGN1 are upregulated in cortical neurons that exhibit resilience to AD, potentially reflecting compensatory responses (*23*). In mice, Nrxn knockout causes deficits in synaptic transmission, plasticity, and cognitive function (*17, 24, 25*). These findings support the idea that protecting Nrxns from AβOs could shift synapses toward tolerance and mitigate cognitive decline. Nrxns therefore represent a critical molecular interface between Aβ pathology and synaptic pathology.

The protein sorting receptor SorCS1 has emerged as a promising candidate for protecting Nrxns against Aβ pathology (*26*). Our previous *in vitro* study has demonstrated that the extracellular domain (ECD) of SorCS1 interacts with β-isoforms of Nrxn (β-Nrxns), competitively blocking AβO-β-Nrxn binding. Notably, SorCS1 rescues AβO-induced impairment of Nrxn-mediated excitatory synapse organization and AβO-induced synaptic toxicity in cultured neurons (*26*). Moreover, SorCS1 promotes axonal trafficking of β-Nrxns and dendritic trafficking of AMPA-type glutamate receptors (AMPARs) (*27*), supporting synaptic integrity. Beyond synaptic roles, previous *in vitro* studies have shown that SorCS1 regulates APP trafficking and suppresses γ-secretase activity, reducing Aβ production (*28–32*). These dual modes of action suggest that SorCS1 protects against both synaptic and amyloid pathology. However, despite strong human genetic evidence linking *SORCS1* to AD (*33–37*) and human single-cell transcriptomic evidence linking *SORCS1* to cognitive resilience in individuals with AD pathology (*38*), its therapeutic potential has not been tested *in vivo*.

In this study, we examined the *in vivo* therapeutic role of SorCS1 in AD by generating an AD mouse model with neuronal SorCS1 overexpression. SorCS1 did not alter Aβ production or plaque deposition but rather ameliorated synaptic pathology and attenuated working memory deficits in AD mice. These effects were associated with restored synaptic expression of β-Nrxns and reduced tau hyperphosphorylation, likely via interactions with GSK3β, a tau kinase. Thus, SorCS1 preserves synaptic integrity and cognitive performance despite ongoing amyloid pathology, positioning it as an “AD resilience-promoting factor” with therapeutic potential for modulation of synaptic connectivity and phosphorylation signaling.

## Results

### Generation of 5xFAD AD mice with neuronal SorCS1 overexpression

SorCS1 is expressed in multiple isoforms (*28*), and overexpression (OE) of the SorCS1b isoform has been shown to attenuate AβO-induced synaptic toxicity in cultured neurons (*26*). To explore the therapeutic potential of SorCS1 in AD model mice, we generated an inducible SorCS1b transgene knock-in (KI) mouse line (herein called SorCS1KI) (**Fig. 1A**). This line overexpresses mouse SorCS1b and tdTomato under the control of the CAG promoter in a Cre recombinase-dependent manner (**Fig. 1A**). The SorCS1KI mice were crossed with 5xFAD mice, a transgenic AD mouse model that carries the transgenes expressing AD-linked mutated human APP and presenilin-1, consequently overproducing Aβ (*39*), as well as with Camk2a-CreERT2 mice for forebrain neuron-specific Cre-mediated recombination upon tamoxifen treatment (*40*). Four experimental mouse groups were established **(Fig. 1B)**: Groups 1 and 2 consist of non-AD mice (wild-type (WT) in the 5xFAD line; 5xFAD^WT/WT^) without (Group 1) or with (Group 2) SorCS1 OE (5xFAD^WT/WT^/Camk2a-CreERT2^+^/SorCS1^KI/KI^ mice with vehicle or tamoxifen), and Groups 3 and 4 consist of AD mice (homozygous 5xFAD; 5xFAD^Tg/Tg^) without (Group 3) or with (Group 4) SorCS1 OE (5xFAD^Tg/Tg^/Camk2a-CreERT2^+^/SorCS1^KI/KI^ mice with vehicle or tamoxifen). SorCS1 OE was induced by oral gavage of tamoxifen for five consecutive days at 1.5 months old. SorCS1 OE was only present in Groups 2 and 4, with the level increased 4-5-fold compared to Group 1 at 6 months old (**Fig. 1C, D**). For the experiments described below, we analyzed the mice using behavioral assays when they were 6 months old followed by the other assays at 6.5-7 months old (**Fig. 1E**).

**Figure 1.**
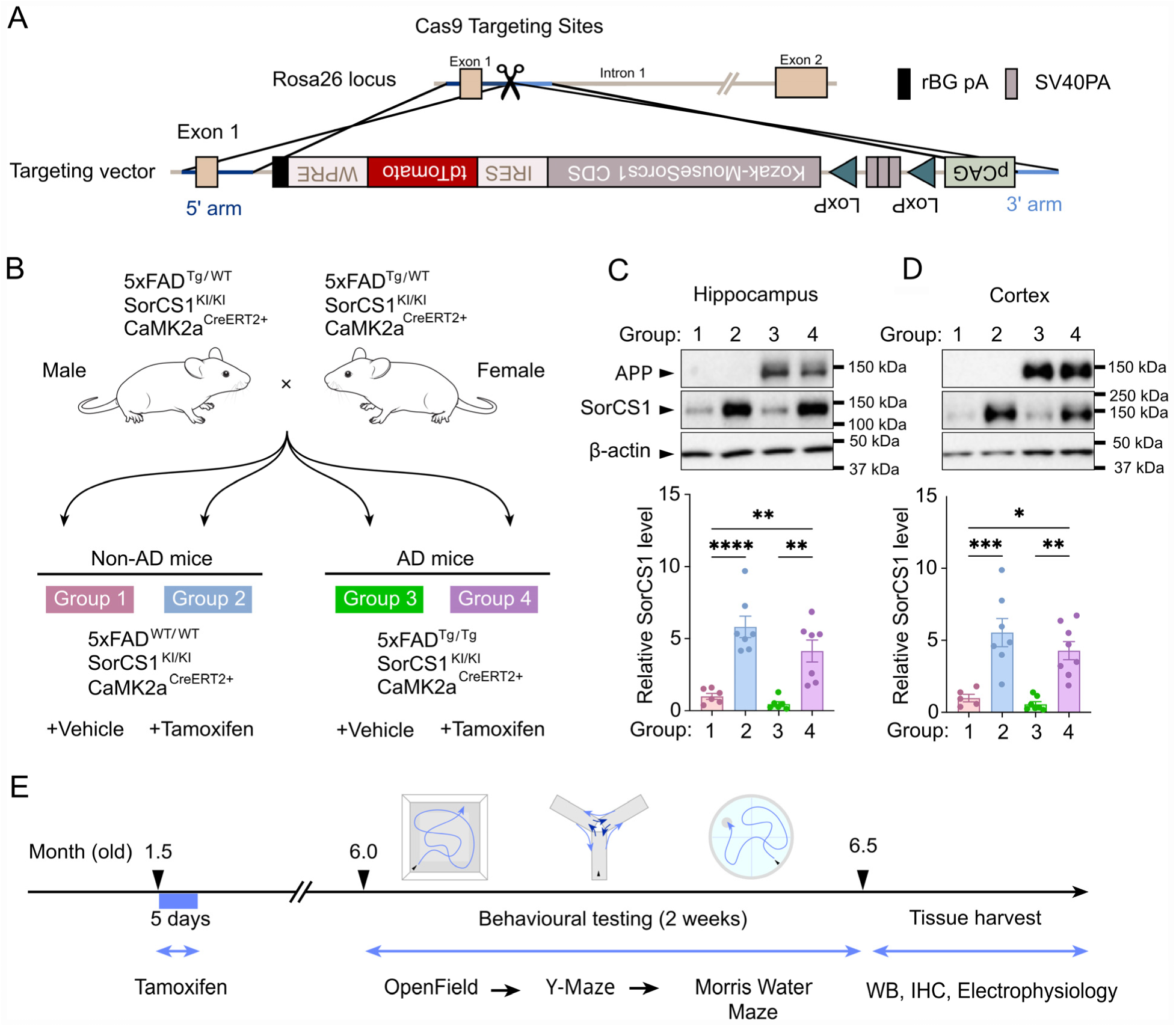
Generation of an inducible SorCS1 knock-in mouse line, validation of experimental cohorts, and study design. **(A)** Schematic representation of the CRISPR/Cas9 knock-in strategy used to insert a CAG-LSL-SorCS1b-tdTomato cassette into the Rosa26 locus. Expression of the transgene remains Cre-dependent until tamoxifen-induced activation in Camk2a-CreERT2 mice, enabling inducible overexpression (OE) of SorCS1b specifically in forebrain excitatory neurons. **(B)** Experimental groups used in this study. Group 1 (Non-AD without SorCS1 OE): 5xFAD^WT/WT^/Camk2a-CreERT2^+^/SorCS1^KI/KI^ mice treated with vehicle; Group 2 (non-AD with SorCS1 OE): 5xFAD^WT/WT^/Camk2a-CreERT2^+^/SorCS1^KI/KI^ mice treated with tamoxifen; Group 3 (AD without SorCS1 OE): 5xFAD^Tg/Tg^/Camk2a-CreERT2^+^/SorCS1^KI/KI^ mice treated with vehicle and Group 4 (AD with SorCS1 OE): 5xFAD^Tg/Tg^/Camk2a-CreERT2^+^/SorCS1^KI/KI^ mice treated with tamoxifen. **(C-D)** Representative western blots and quantification of SorCS1 and APP protein levels in hippocampal (**C**) and cortical (**D**) lysates from the indicated groups. Protein expression was normalized to β-actin and expressed relative to the mean value of Group 1. Data are presented as mean ± SEM. Statistical significance was determined by two-way ANOVA followed by Sidak’s multiple comparisons test. *P < 0.05, **P < 0.01, \*\*\**P* < 0.001, ****P < 0.0001; ns, not significant. (**E**) Experimental timeline. Tamoxifen was administered once daily for 5 consecutive days beginning at 1.5 months of age to induce SorCS1 OE. Behavioral testing, including the open-field test, Y-maze, and Morris water maze, was performed at 6 months of age over a 2-week period. Mice were euthanized at 6.5 months of age to collect brains for biochemical and histological analyses.

### Neuronal SorCS1 overexpression prevents short-term working memory deficits in female 5xFAD mice

5xFAD mice typically display impairments in learning and memory by 4-5 months of age (*41, 42*). To assess spatial working memory of mice in Groups 1-4, we performed a Y-maze alternation test (*43*) (**Fig. 2A-E**). Male AD mice without SorCS1 OE (Group 3) did not show significant reduction of alternation percentage or total arm entry compared with non-AD males (Group 1)(**Fig. 2A, D, E**), indicating that 5xFAD males in our cohorts did not display pronounced working memory deficits at 6 months old. In contrast, in females, the spontaneous alternation percentage of AD mice without SorCS1 OE (Group 3) was lower than that of non-AD mice without SorCS1 OE (Group 1) (**Fig. 2A, B**), consistent with previous studies showing impaired working memory in 5xFAD mice (*39, 42*). Notably, AD females with SorCS1 OE (Group 4) showed a higher alternation percentage than AD females without OE (Group 3), reaching levels comparable to non-AD controls (Group 1) (**Fig. 2A, B**). This indicates that neuronal SorCS1 OE in the forebrain prevents working memory deficits in female 5xFAD mice. Moreover, non-AD females with SorCS1 OE (Group 2) showed alternation percentages at a similar level to non-AD females without SorCS1 OE (Group 1), indicating that SorCS1 OE does not affect working memory in non-AD healthy condition (**Fig. 2A, B**). Female mice across all groups exhibited comparable numbers of total arm entries (**Fig. 2C**), confirming that differences in alternation percentage were not due to changes in overall locomotor activity.

**Figure 2.**
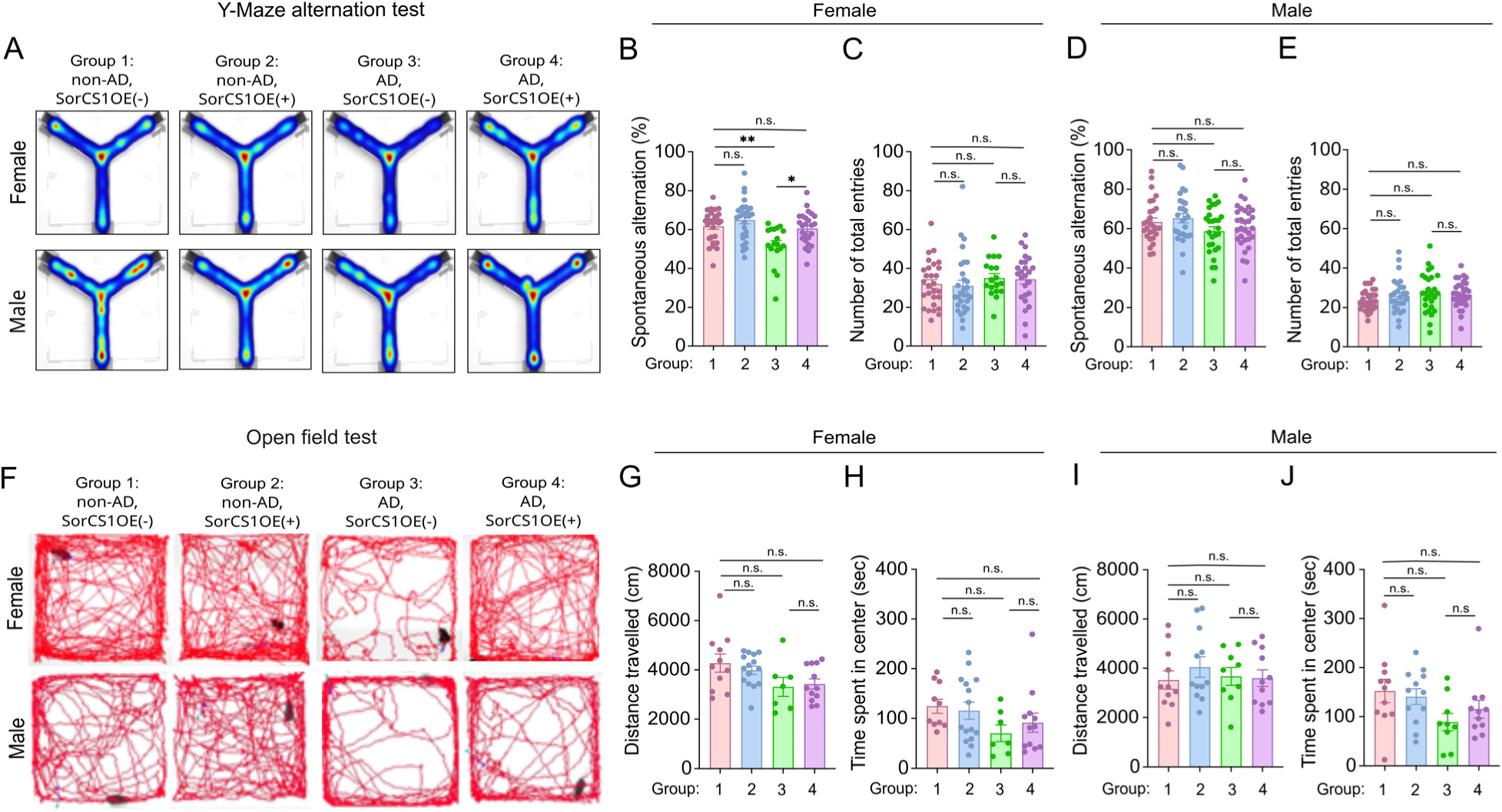
Neuronal SorCS1 overexpression attenuates short*-*term working memory deficits in female 5xFAD mice. (**A**) Representative Y-maze heat maps showing exploration patterns of female and male mice from Groups 1–4. (**B**–**E**) Quantification of Y-maze performance showing the percentage of spontaneous alternation (**B**, **D**) and the total number of arm entries (**C, E**) in female (**B**, **C**) and male (**D**, **E**) mice. Group sizes were: Group 1 (non-AD, without SorCS1 OE), *n* = 28 females and 29 males; Group 2 (non-AD with SorCS1OE), *n* = 29 females and 27 males; Group 3 (AD without SorCS1OE), *n* = 18 females and 27 males; and Group 4 (AD with SorCS1OE), *n* = 25 females and 31 males. (**F**) Representative open-field tracking traces illustrating locomotion of female and male mice from Groups 1-4. (**G**–**J**) Quantification of open-field behavior showing total distance traveled (**G**, **I**) and time spent in the center zone (**H**, **J**) in female (**G**, **H**) and male (**I**, **J**) mice. Group sizes were: Group 1, *n* = 11 females and 11 males; Group 2, *n* = 15 females and 12 males; Group 3, *n* = 7 females and 9 males; and Group 4, *n* = 12 females and 11 males. Data are presented as mean ± SEM. Statistical analyses were performed using two-way ANOVA followed by Sidak’s multiple comparisons test. *P < 0.05, **P < 0.01, \*\*\**P* < 0.001, ****P < 0.0001; ns, not significant.

Next, we conducted an open field test to examine locomotor activity and anxiety levels in Groups 1-4 (**Fig. 2F-J**). In both sexes, mice in all groups exhibited comparable total distances traveled (**Fig. 2G,I**), indicating intact locomotor activity across all groups and consistent with the total arm entry results of the Y-maze tests. To assess anxiety, the time spent in the center of the arena was measured (**Fig. 2H, J**). Although Group 3 mice of both sexes tended to spend less time in the center compared to Group 1 mice, no statistical significance was detected in any group comparisons (**Fig. 2H, J**). Together, these data suggest that locomotor activity and anxiety levels are normal across the groups in both sexes.

Hallmark impairments in 5xFAD mice are deficits in spatial learning and long-term reference memory (*41, 44*). To assess these impairments, we conducted Morris water maze testing (*45*) (**Supplementary Fig. 1**). In both sexes, mice from Groups 1 and 2 showed comparable escape latencies that progressively declined over the five training days (**Supplementary Fig. 1C, F**), indicating intact spatial learning. In contrast, mice from Groups 3 and 4 required significantly longer escape times than Groups 1 and 2 (**Supplementary Fig. 1C, F**), although Group 4 mice tended to reach the hidden platform within 60-sec test period more frequently than those in Group 3 (83.3% (9 mice out of 11 mice) in Group 4 but 50% (6 mice out of 12 mice) in Group 3, P = 0.1095 by Chi-square test). In the probe test, Group 3 and 4 mice performed significantly fewer platform crossings and spent less time in the target quadrant compared to mice from Groups 1 and 2 (**Supplementary Fig. 1I, J**). Moreover, a swimming velocity assessment revealed that both male and female mice in Groups 3 and 4 swam significantly slower than those in Groups 1 and 2 across all training sessions (**Supplementary Fig. 1D, G**). These results could suggest that neuronal SorCS1 OE in the forebrain may not prevent the deficits in spatial learning, reference memory or skilled motor function in AD mice. However, as previous studies have noted(*45, 46*), motor-related impairments such as reduced swimming speed and increased thigmotaxis confound accurate assessment of spatial learning and reference memory. Consequently, the Morris water maze is likely too demanding to serve as a reliable measure of long-term learning and memory for the 5xFAD AD mice in our cohorts. Given the differences observed in females in the Y-maze tests (**Fig. 2B**), subsequent analyses in this study focused on females.

### Neuronal SorCS1 overexpression restores impaired basal excitatory synaptic transmission in the hippocampus of 5xFAD hippocampus

To examine whether the protective role of SorCS1 relates to synaptic function, we measured field excitatory postsynaptic potentials (fEPSPs) evoked by stimulation of Schaffer collateral (SC) afferents in the dorsal CA1 hippocampus of all female groups (**Fig. 3A, B**). Analysis of input-output (I-O) responses of SC-CA1 synapses, defined by the relationship between the fEPSP slope and the fiber volley (FV) amplitude, revealed that AD mice without SorCS1 OE (Group 3) exhibited a significant reduction in I-O responses compared with non-AD controls (Group 1) (**Fig. 3B**). Importantly, the I-O relationships of AD mice with SorCS1 OE (Group 4) and non-AD mice with SorCS1 OE (Group 2) were indistinguishable from those of non-AD controls (Group 1) (**Fig. 3B**). These results indicate that neuronal SorCS1 OE restores basal excitatory synaptic transmission deficits in AD mice, while having no significant effect in non-AD conditions.

**Figure 3.**
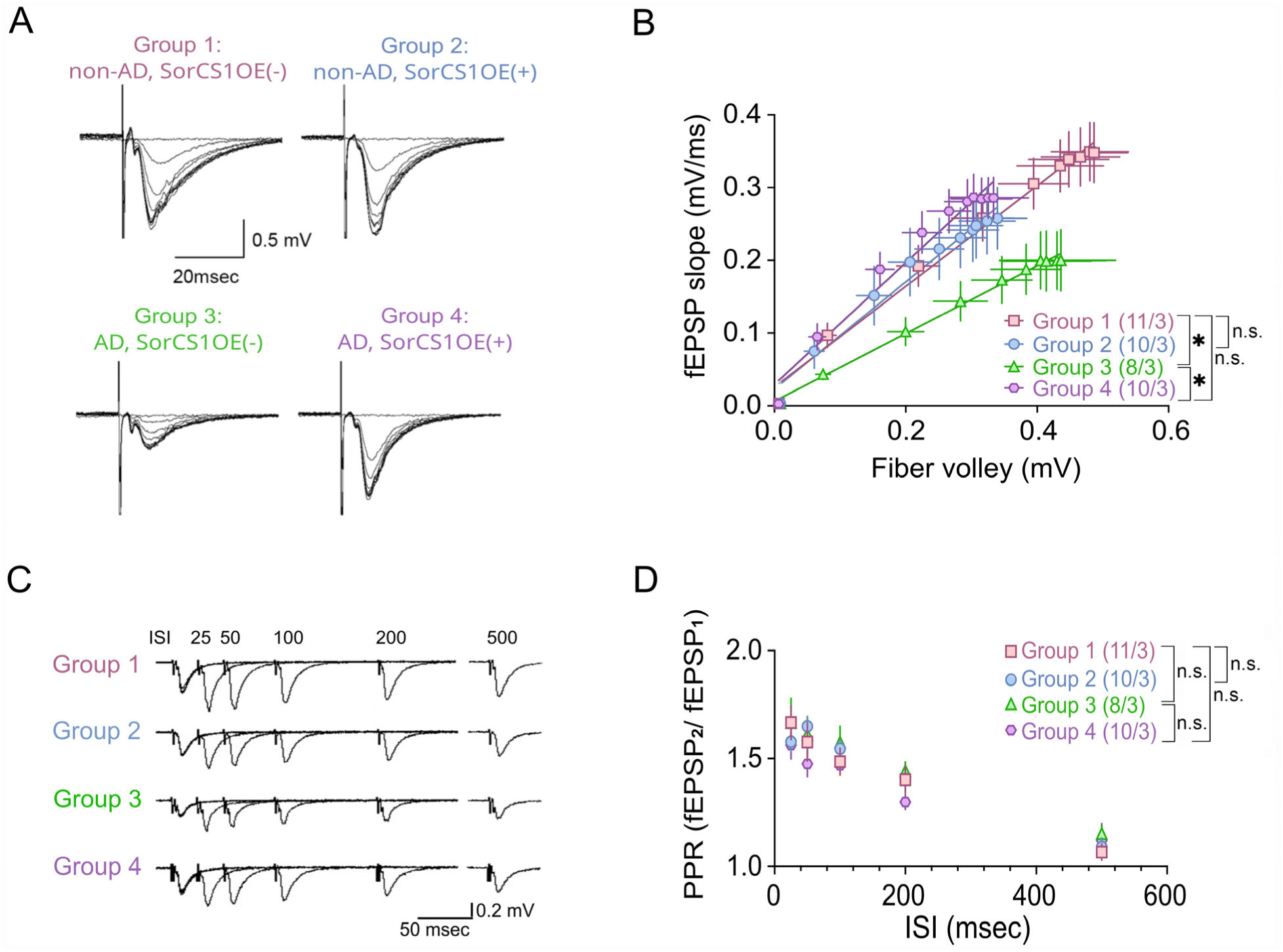
Neuronal SorCS1 overexpression restores impaired basal synaptic transmission but does not alter paired-pulse facilitation in the hippocampus of 5xFAD females. **(A)** Representative field excitatory postsynaptic potential (fEPSP) traces recorded from the hippocampal CA1 stratum radiatum of female mice in Group 1-4 following stepwise increases in stimulation intensity. Scale bar, 0.5 mV, 20 ms. **(B)** Input–output (I/O) curves showing the relationship between fiber volley (FV) amplitude and fEPSP slope in the indicated groups. Basal synaptic transmission was assessed by linear regression analysis of FV-fEPSP relationships. *P < 0.05, **P < 0.01, ***P < 0.001, ****P < 0.0001; ns, not significant. **(C)** Representative paired-pulse facilitation (PPF) traces recorded at interstimulus intervals (ISIs) of 25, 50, 100, 200, and 500 ms from hippocampal CA1 synapses in Groups 1-4. **(D)** Quantification of paired-pulse ratios (PPR; fEPSP₂/fEPSP₁) across the indicated ISIs in each group. PPR data were analyzed using two-way repeated-measures ANOVA. Data are presented as mean ± SEM. Group sizes: n = 11, 10, 8, and 10 hippocampal slices obtained from 3 female mice per group for Groups 1, 2, 3, and 4, respectively.

We next examined paired-pulse facilitation (PPF), a form of presynaptic short-term plasticity(*47*), at the SC-CA1 synapses across all groups (**Fig. 3C, D**). PPF was comparable among all groups at multiple inter-stimulation intervals ranging from 25 to 500 msec, indicating that SC-CA1 synapses in 5xFAD mice exhibit no significant impairment in glutamate release probability (Pr) and that SorCS1 OE does not alter Pr in either non-AD or AD conditions (**Fig. 3C, D**). Altogether, these results suggest that SorCS1-mediated restoration of synapse function in AD mice results from the preservation of functional synapses and/or compensatory enhancement of postsynaptic function.

### Neuronal SorCS1 overexpression preserves hippocampal synaptic integrity in 5xFAD mice

We next investigated synaptic integrity in the mouse groups. Our recent *in vitro* study demonstrated that SorCS1 OE rescues excitatory synapse loss and postsynaptic density (PSD) shrinkage induced by AβO treatment in cultured hippocampal neurons (*26*), suggesting that SorCS1 counteracts Aβ-induced synaptic damage. To evaluate this effect *in vivo*, we performed western blot analyses on hippocampal synaptosome samples from all female groups to examine the synaptic expression of PSD95 (an excitatory postsynaptic scaffold protein) and synaptophysin (SYP; a presynaptic vesicle protein) (**Fig. 4A-D**). As expected, synaptic levels of both proteins were reduced in AD mice without SorCS1 OE (Group 3) compared to non-AD controls (Group 1) (**Fig. 4C, D**). Notably, AD mice with SorCS1 OE (Group 4) exhibited significantly higher synaptic expression of PSD95 and SYP than Group 3 mice (**Fig. 4C, D**). These findings indicate that SorCS1 OE restores PSD95 and SYP expression in 5xFAD synapses.

**Figure 4.**
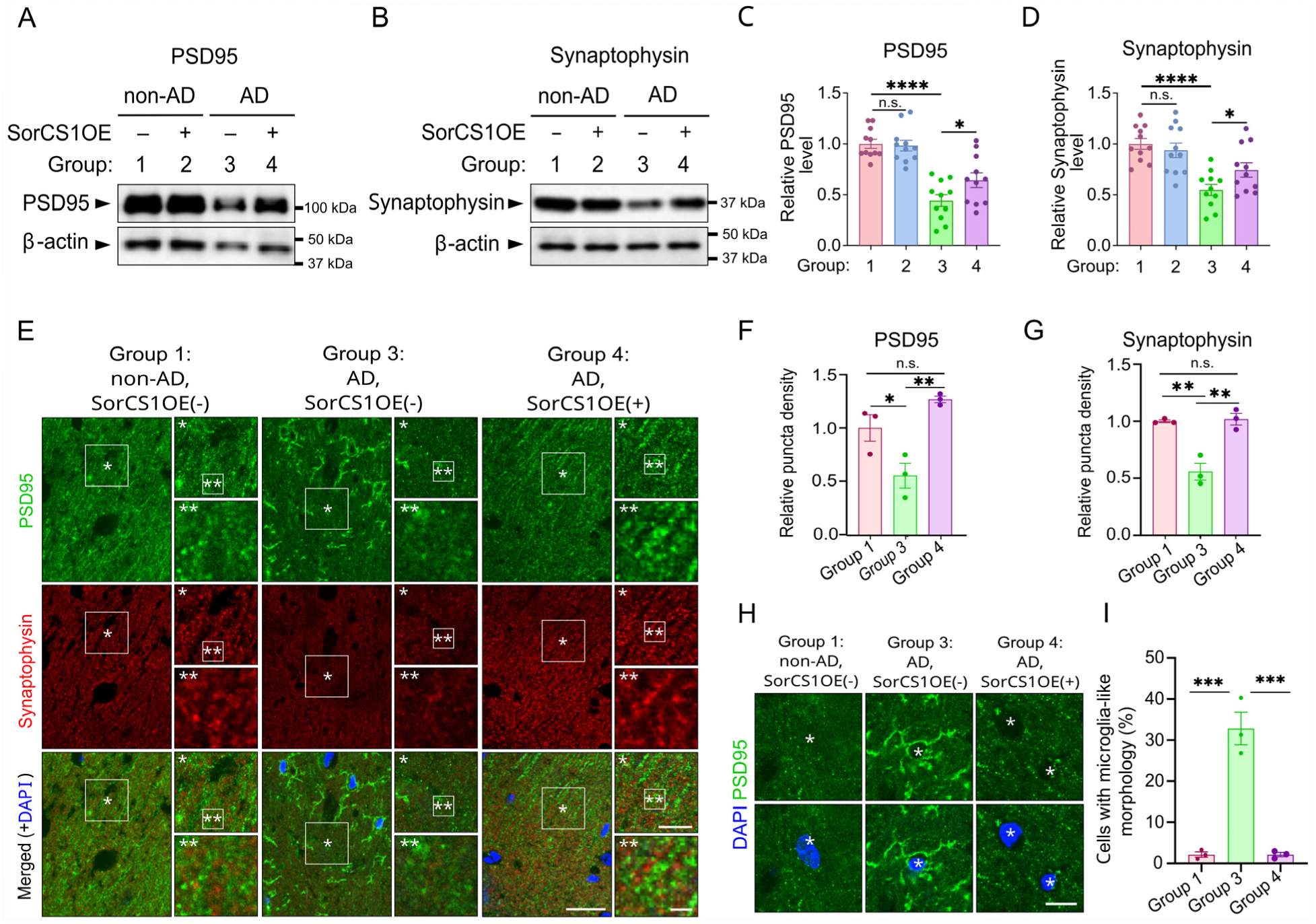
Neuronal SorCS1 overexpression restores impaired excitatory synapse integrity in the hippocampus of female 5xFAD mice. (**A-D)** Representative immunoblots **(A, B)** and corresponding quantification **(C, D)** of PSD95 and synaptophysin in hippocampal crude synaptosomal fractions isolated from female mice in Groups 1-4. Protein expression was normalized to β-actin and expressed relative to the mean value of Group 1. Statistical analyses were performed using two-way ANOVA followed by Sidak’s multiple comparison test. Group sizes: n = 11 female mice per group. **(E)** Representative confocal images of PSD95 (green) and synaptophysin (red) immunostaining in the CA1 stratum radiatum of the hippocampus with DAPI nuclear counterstaining (blue). Insets show medium- and high-magnification views of the boxed regions. Scale bars: 25 μm (low magnification), 5 μm (medium magnification), and 1.5 μm (high magnification). (**F**, **G**) Quantification of PSD-95 (**F**) and synaptophysin (**G**) puncta density. Protein expression was normalized to β-actin and expressed relative to the mean value of Group 1. Statistical analyses were performed using two-way ANOVA followed by Sidak’s multiple comparison test. Group sizes: n = 3 female mice per group. For each mouse, 12 images were analyzed and averaged to obtain a single value for statistical comparison. Scale bar, 25 μm. (**H, I**) Representative images (**H**) and quantification (**I**) of PSD95-positive cells exhibiting microglia-like morphology. Statistical analyses were performed using two-way ANOVA followed by Sidak’s multiple comparison test. Group sizes: n = 3 female mice per group. For each mouse, 12 images were analyzed and averaged to obtain a single value for statistical comparison. Data are presented as mean ± SEM. Statistical significance is indicated as follows: *P < 0.05, **P < 0.01, ***P < 0.001, ****P < 0.0001; ns, not significant.

We then performed immunohistochemistry to assess PSD95 and SYP immunoreactivity in the stratum radiatum of the CA1 dorsal hippocampus of Group 1, 3 and 4 females (**Fig. 4E-I**). In non-AD mice (Group 1), numerous small PSD95 and SYP puncta, corresponding to synaptic structures, were detected (**Fig. 4E, left**). In contrast, AD mice without SorCS1 OE (Group3) showed a remarkable reduction in synaptic puncta of both proteins (**Fig. 4E middle**). Notably, AD mice with SorCS1 OE (Group 4) displayed abundant synaptic PSD95 and SYP puncta, resembling the pattern in non-AD controls (Group 1) (**Fig. 4E right**). Quantitative analysis confirmed that puncta density of PSD95 and SYP were significantly reduced in Group 3 compared with Group1, whereas Group 4 values were significantly elevated relative to Group 3 and comparable to Group 1 values (**Fig. 4F, G**). These results corroborate the synaptosome western blot findings (**Fig. 4A-D**) and are further supportive of protective effects of SorCS1 on Aβ-induced synaptic deficits and contribution of neuronal SorCS1 to preserving synapse integrity in 5xFAD mice.

Unexpectedly, PSD95 immunohistochemistry also revealed numerous microglia-like immunoreactive structures in Group 3 but not in Group 1 or Group 4 (**Fig. 4H, I**). Given that activated microglia upregulate PSD95(*48*), this observation suggests that microglial activation occurs in AD mice and that neuronal SorCS1 OE may suppress microglial activation in the hippocampus.

### Neuronal SorCS1 OE restores synaptic expression of *β*-Nrxn, but not *α*-Nrxn, in 5xFAD mice

In healthy conditions, SorCS1 facilitates axonal targeting of β-Nrxn through their extracellular interaction (*27*) and of α-Nrxn through intracellular mechanisms (*49*). Under pathological conditions, SorCS1 competitively inhibits extracellular AβO binding to β-Nrxn, ameliorating AβO-induced synaptic defects in cultured hippocampal neurons (*26*). Given these findings, we examined synaptic expression of α- and β-Nrxn in synaptosome preparations from all female groups (**Fig. 5**). In both hippocampal and cortical synaptosomes, AD mice without SorCS1 OE (Group 3) showed significantly reduced levels of α- and β-Nrxn compared with non-AD controls (Group 1) (**Fig. 5A-F**). In contrast, AD mice with SorCS1 OE (Group 4) exhibited significantly higher expression of β-Nrxn, but not α-Nrxn, relative to Group 3 (**Fig. 5A-F**). Thus, SorCS1 OE selectively rescues synaptic expression of β-Nrxns in AD synapses. Given our previous findings that AβO-β-Nrxn interactions reduce β-Nrxn expression on the axon surface (*19*) and that SorCS1 competitively blocks this interaction (*26*), these results suggest that SorCS1 preserves synaptic transmission and integrity in AD mice by protecting β-Nrxns from AβO binding.

**Figure 5.**
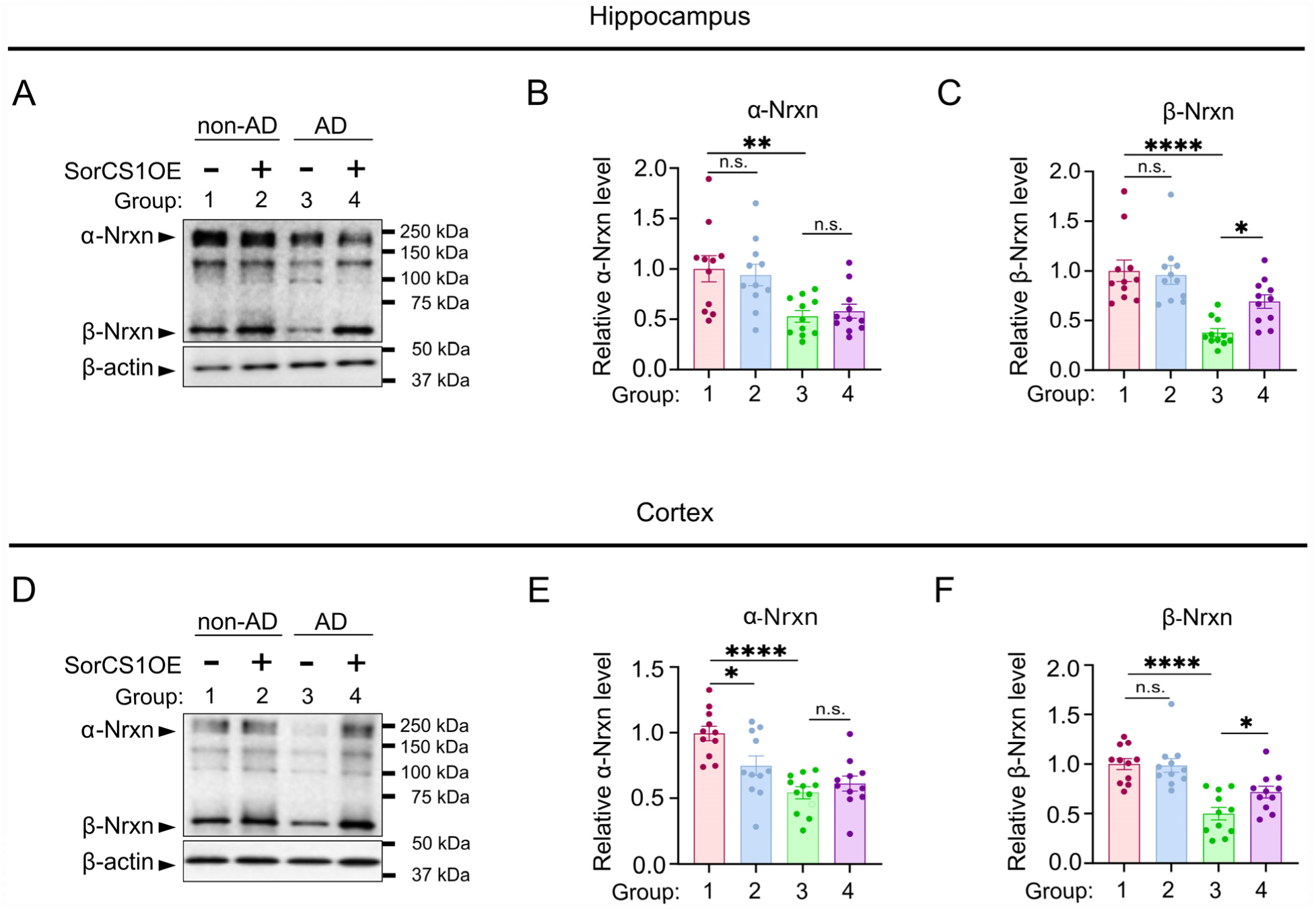
Neuronal SorCS1 overexpression restores synaptosomal *β*-neurexin levels in the hippocampus and cortex. (**A, D**) Representative immunoblots of α-neurexin (α-Nrxn) and β-neurexin (β-Nrxn) in hippocampal (**A**) and cortical (**D**) synaptosomal fractions isolated from female mice in Groups 1-4. Proteins were detected using a pan-neurexin antibody (ABN161-I). (**B, C, E, F**) Quantification of α-Nrxn (**B**, **E**) and β-Nrxn (**C**, **F**) protein levels in hippocampal (**B**, **C**) and cortical (**E**, **F**) synaptosomal fractions from Groups 1-4. Protein expression was normalized to β-actin and expressed relative to the mean value of Group 1. Data are presented as mean ± SEM. Statistical analyses were performed using two-way ANOVA followed by Sidak’s multiple comparisons test. Group sizes: n = 11 mice per group. *P < 0.05, **P < 0.01, ***P < 0.001, ****P < 0.0001; ns, not significant.

Given that SorCS1 also regulates dendritic trafficking of AMPARs in cortical neurons (*27*), we next investigated synaptic expression of the AMPAR subunits GluA1 and GluA2 in cortical synaptosomes (**Supplementary Fig. 2**). AD mice without SorCS1 OE (Group 3) displayed significantly lower levels of both GluA1 and GluA2 compared with non-AD controls (Group 1). In contrast, AD mice with SorCS1 OE (Group 4) exhibited significantly increased expression of both AMPAR subunits relative to Group 3. However, in hippocampal synaptosomes, SorCS1 OE failed to rescue the AD-associated reductions in GluA1 and GluA2 expression. These results indicate that the rescue effect of SorCS1 on synaptic AMPAR expression is region-specific, enhancing AMPAR levels in the cortex but not in the hippocampus.

### Neuronal SorCS1 overexpression has no effect on amyloid pathology in 5xFAD mice

Previous *in vitro* studies have demonstrated that SorCS1 OE suppresses Aβ production (*30, 31*). To determine whether SorCS1 OE exerts similar effects *in vivo*, we investigated amyloid plaque deposition and AβO expression in AD female mice with or without SorCS1 OE (**Fig. 6**). Contrary to the previous *in vitro* findings, our immunohistochemistry showed comparable amyloid plaque densities between Group 3 and Group 4 in the hippocampal CA1, dentate gyrus, and cortical regions (**Fig. 6A-D**). Consistently, our immunoblot analysis revealed no significant differences in AβO (tetramer) abundance between these groups in the hippocampal and cortical synaptosomes (**Fig. 6E, F**). Although female 5xFAD mice typically exhibit more severe phenotypes than males (*50*), we also investigated male mice, again finding no significant difference between Group 3 and Group 4 in amyloid deposition and AβO abundance (**Supplementary Fig. 3**). Thus, despite its protective effects on preserving synapse function and integrity under AD conditions (**Figs. 3-5**), SorCS1 OE does not mitigate *in vivo* amyloid pathology. These findings suggest that synapse pathology and cognitive impairment can be alleviated even in the presence of ongoing Aβ accumulation, supporting the concept of cognitive resilience to AD (*12, 13*).

**Figure 6.**
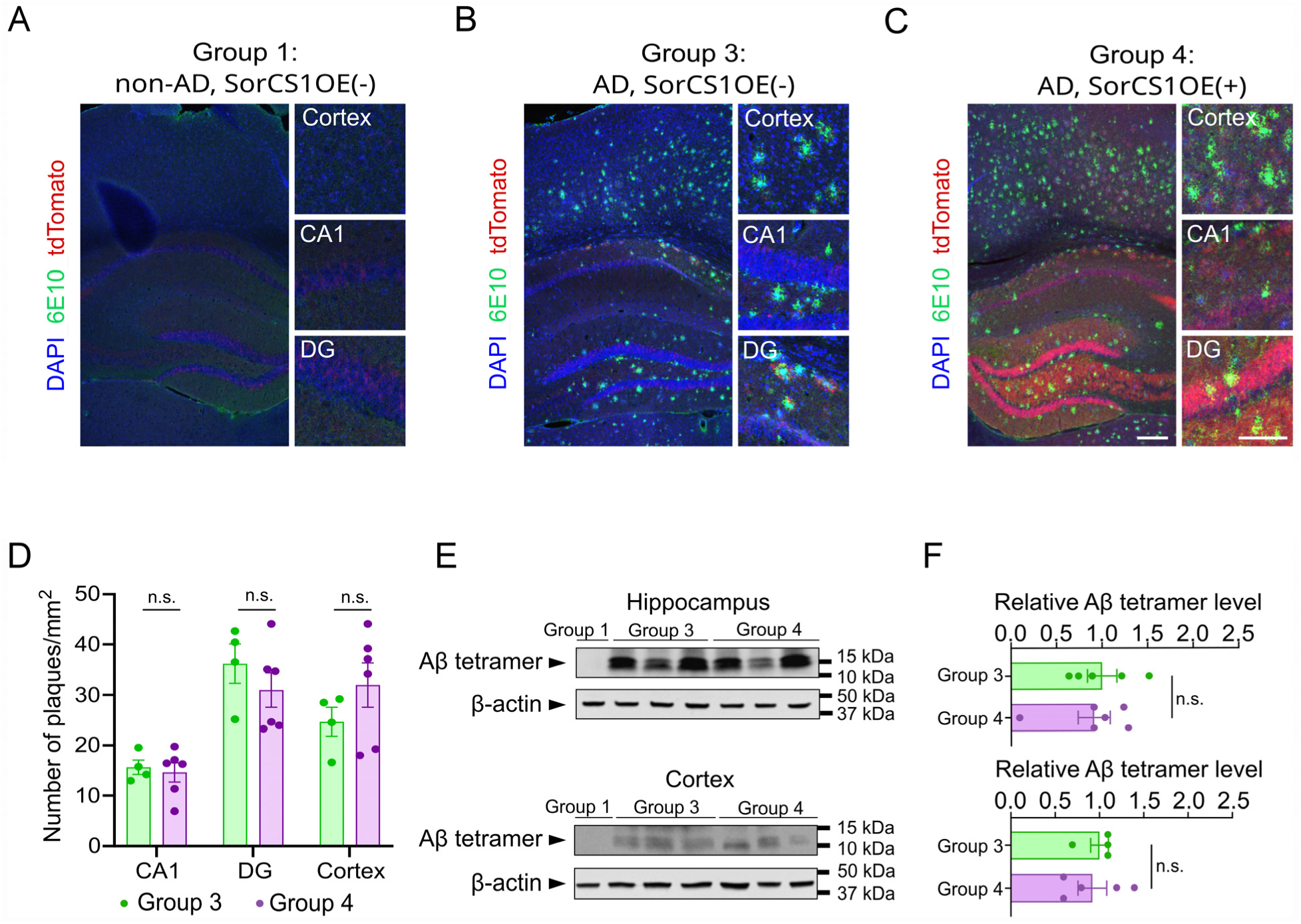
Neuronal SorCS1 overexpression does not affect amyloid plaque burden or A*β* tetramer levels in female 5xFAD mice. (**A-C**) Representative immunofluorescence images of amyloid plaques labeled with 6E10 antibody (green) in female mice from Groups 1, 3 and 4. tdTomato fluorescence, indicating transgene expression, is shown in red, and nuclei were counterstained with DAPI (blue). Higher-magnification images of the hippocampal CA1 region, and dentate gyrus (DG), and cortex are shown on the right. Scale bars: 500 μm (low magnification) and 250 μm (high magnification). (**D**) Quantification of 6E10-positive plaque density (plaques/mm²) in the CA1, DG and cortex of female mice from Groups 3 and 4. Group sizes: n = 4 and 6 mice for Group 3 and Group 4, respectively. ns, not significant. (**E, F**) Representative immunoblots (**E**) and quantification (**F**) of Aβ tetramers in hippocampal (top) and cortical (bottom) lysates from Group 3 and 4 female mice. Aβ tetramer expression was normalized to β-actin and expressed relative to the mean value of Group 3. Group sizes: n = 5 and 6 mice for Groups 3 and 4, respectively, in the hippocampus, and n = 4 and 5 mice for Groups 3 and 4, respectively, in the cortex. ns: not significant. Data represent mean ± SEM. Statistical analyses were performed using unpaired two-tailed Student’s *t*-tests.

### Neuronal SorCS1 overexpression mitigates tau hyperphosphorylation in 5xFAD mice

We next examined the effect of SorCS1 OE on tau pathology in AD mice (**Fig. 7**). Given that tau hyperphosphorylation has been reported in 5xFAD mice (*51*), we assessed phosphorylated tau (p-tau) and total tau levels in hippocampal synaptosomes from each group by western blot analysis (**Fig. 7A-C**). As expected, AD mice without SorCS1 OE (Group 3) exhibited elevated levels of p-tau per total tau (p-tau/tau) compared with non-AD controls (Group 1), with statistically significant differences in hippocampal synaptosomes (**Fig. 7A, B**). Notably, AD mice with SorCS1 OE (Group 4) showed a significantly reduced p-tau/tau ratio compared with Group 3, restoring values to levels comparable with non-AD controls (Group 1) (**Fig. 7A, B**). In contrast, total tau normalized to β-actin (tau/β-actin) was significantly lower in both Group 3 and Group 4 relative to Group 1 (**Fig. 7A, C**). Together, these results suggest that SorCS1 OE attenuates tau hyperphosphorylation in 5xFAD mice, although it does not normalize the reduced total tau levels in synaptosomes.

**Figure 7.**
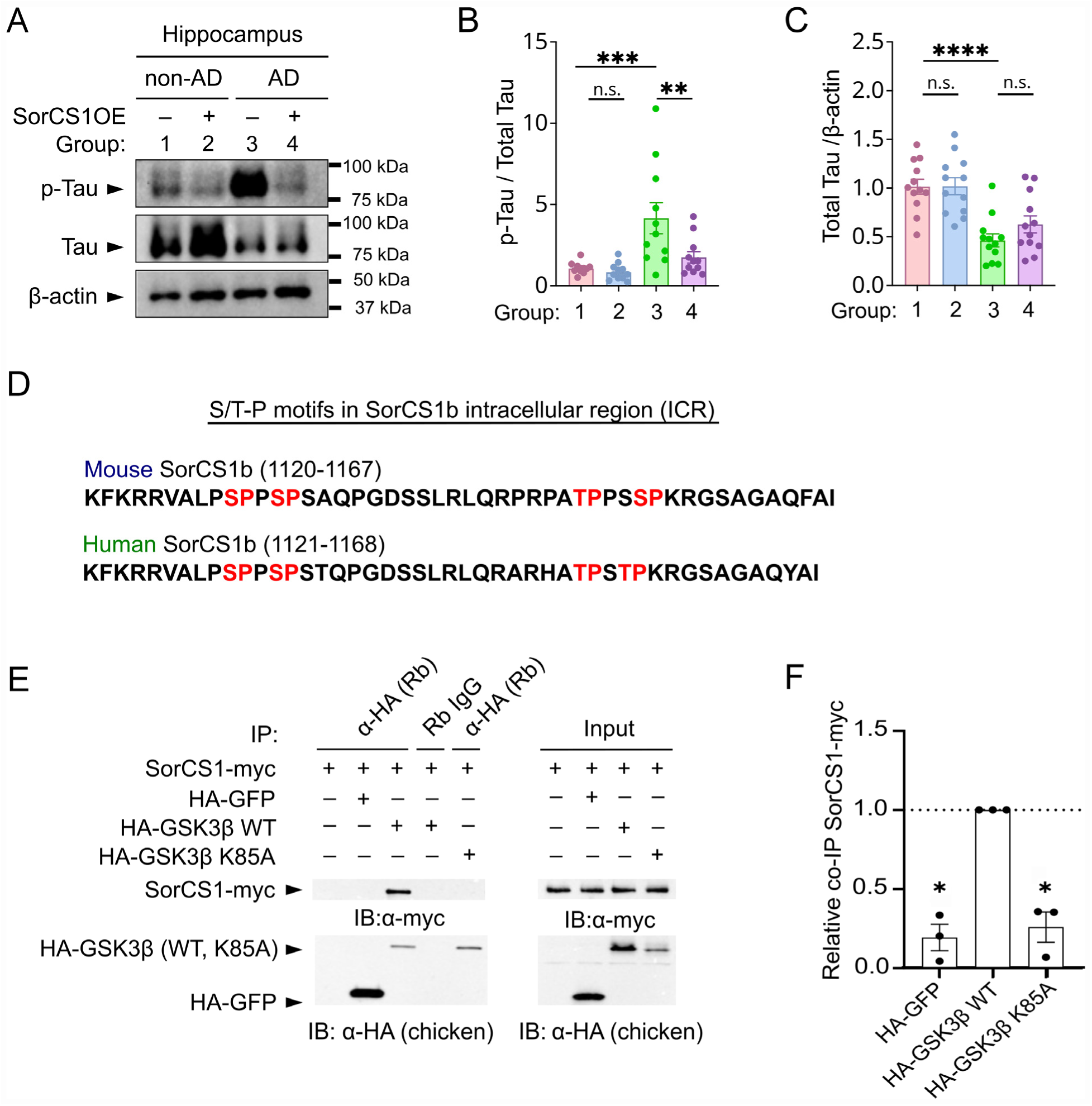
Neuronal SorCS1 overexpression attenuates tau hyperphosphorylation in the hippocampus of female 5xFAD mice, and SorCS1 interacts with the tau kinase GSK3*β* in a kinase activity-dependent manner. (**A**) Representative western blots showing phosphorylated tau (p-Tau), total tau (Tau) and β-actin (loading control) in total hippocampal lysates obtained from female mice in Groups 1-4. (**B**, **C**) Quantification of the p-Tau/Tau ratio **(B)** and the Tau/β-actin ratio **(C)** in hippocampal lysates from female mice in Groups 1-4. Protein expression was normalized to β-actin and expressed relative to the mean value of Group 1. Data are presented as mean ± SEM, with individual mice shown as dots. Statistical analyses were performed using two-way ANOVA followed by Sidak’s multiple comparisons test. Group sizes: n = 11 female mice per group. *P < 0.05, **P < 0.01, ***P < 0.001, ****P < 0.0001; ns, not significant. **(D)** Amino acid sequences of the intracellular region (ICR) of mouse and human SorCS1b (UniProt IDs: Q9JLC4 and Q8WY21, respectively). The ICR contains four serine/threonine-proline (S/T-P) motifs (underlined), which represent consensus phosphorylation sites for proline-directed protein kinases, including the tau kinase GSK3β. **(E)** Co-immunoprecipitation (Co-IP) analysis demonstrates an interaction between SorCS1 and GSK3β that depends on GSK3β kinase activity. HEK293T cells were co-transfected with SorCS1-myc and HA-GSK3β encoding either wild-type (WT) or kinase-dead (K85A). Cell lysates were subjected to immunoprecipitation using a rabbit polyclonal anti-HA antibody, followed by immunoblotting with a mouse monoclonal anti-myc antibody and a chick polyclonal anti-HA antibody. Mouse IgG was used as a negative immunoprecipitation control, and input lysates confirmed comparable expression levels of the transfected proteins. HA-GFP served as a negative control for HA-tagged proteins. **(F)** Quantification of co-immunoprecipitated SorCS1 normalized to the amount of immunoprecipitated HA-tagged protein. SorCS1 levels were expressed relative to the HA-GSK3β WT condition in each experiment. Data are presented as mean ± SEM, with each dot representing one independent culture preparation. Statistical analyses were performed using one sample t-test. n = 3 independent experiments. *P < 0.05.

### SorCS1 interacts with the tau kinase GSK3*β*

The intracellular region (ICR) of mouse and human SorCS1b (Uniprot ID: Q9JLC4 and Q8WY21, respectively) contains four serine/threonine–proline (S/T-P) motifs (**Fig. 7D**), the consensus target sequence for proline-directed protein kinases (PDPKs) (*52*). Several PDPKs, including GSK3β are well known to phosphorylate tau (*53*), raising the possibility that SorCS1 engages PDPKs to modulate tau phosphorylation. To test this, we performed co-immunoprecipitation (co-IP) assays using HEK293T cells. SorCS1b-myc precipitated with the anti-HA antibody (αHA), but not control IgG, only when the cells were co-transfected with HA-tagged GSK3β wild-type construct (HA-GSK3β WT), whereas no co-IP was detected with the kinase-dead variant HA-GSK3β K85A (**Fig. 7E, F**). These findings indicate that SorCS1b interacts with GSK3β in a kinase activity-dependent manner.

## Discussion

In this study, we generated and comprehensively characterized 5xFAD mice with forebrain-specific neuronal overexpression of SorCS1 and demonstrated that SorCS1 confers synaptic and cognitive preservation despite substantial progression of amyloid pathology. Mechanistically, SorCS1-mediated synaptic protection was linked to the restoration of synaptic β-Nrxn expression, offering new molecular insights into its role in maintaining synaptic function and integrity. In addition, SorCS1 attenuated tau hyperphosphorylation in 5xFAD hippocampus, likely through interactions with GSK3β. Together, these findings uncover an unrecognized function of this protein-sorting receptor in regulating synapse stability and tau phosphorylation in AD, establishing SorCS1 as a novel AD resilience-promoting factor with multifaceted molecular mechanisms.

A striking and unexpected finding of this study is that neuronal SorCS1 OE preserves synaptic integrity, basal transmission, and working memory despite failing to mitigate amyloid pathologies such as plaque deposition or AβO expression. This result supports the notion that directly targeting Aβ peptides or amyloid cascades may not be essential for maintaining cognitive function, aligning with the concept of cognitive resilience in AD (*12–15*) and the failure of many clinical trials that target Aβ peptides and amyloid cascades (*6*). Instead, SorCS1 emerges as a key molecule that promotes synaptic resilience and cognitive preservation during the progression of amyloid pathology. In contrast to our *in vivo* results, previous *in vitro* studies using HEK293T cells transiently transfected with APP reported that SorCS1 OE reduces Aβ production (*30*). This discrepancy may reflect differences in experimental contexts and/or SorCS1 isoform usage. In our study, we examined the SorCS1b isoform, which is preferentially targeted to the plasma membrane (*32*), whereas the previous study investigated SorCS1c, an isoform enriched in endosomes but not at the plasma membrane(*32*). Future work should address whether SorCS1c OE mitigates amyloid pathology under *in vivo* conditions.

SorCS1 acts as a key regulator of protein sorting under physiological conditions, promoting the trafficking of both β- and α-Nrxns to the axonal surface (*27, 49*). SorCS1 directly interacts with β-Nrxns, via its ECD (*27*), which binds the N-terminal histidine-rich domain (HRD) of β-Nrxns (*26*). In contrast, the SorCS1 ECD does not bind α-Nrxns (*26*); instead, its intracellular domain (ICD) recruits the Rab effector Rip11 to mediate α-Nrxn trafficking (*49*). Thus, SorCS1 orchestrates axonal trafficking of β- and α-Nrxns through distinct mechanisms. Under pathological conditions associated with Aβ, SorCS1 exerts an additional role: binding of its ECD to β-Nrxns competitively inhibits the AβO-β-Nrxn interaction, which also depends on the β-Nrxn HRD (*26*). These findings suggest that SorCS1-dependent restoration of synaptic β-Nrxn expression in 5xFAD mouse brains may rely on this extracellular interaction, highlighting the SorCS1 ECD and the β-Nrxn HRD as potential therapeutic targets for synapse preservation in AD. Notably, SorCS1 does not restore synaptic α-Nrxn expression in 5xFAD brains, underscoring again the critical role of its extracellular interaction with β-Nrxns. In this study, we focused on SorCS1b, while Rip11 interaction has been reported for SorCS1c (*49*), potentially reflecting isoform-specific functions.

Another unexpected finding of this study is that SorCS1 OE mitigates tau hyperphosphorylation despite no previous evidence linking SorCS1 to the regulation of protein phosphorylation. We demonstrate interactions between SorCS1b and GSK3β, although whether and how this interaction affects GSK3β kinase activity remains unresolved. Notably, PhosphositePlus, an online catalogue of experimentally observed post-translational modifications (*54*), reports phosphorylation of several serine/threonine (S/T) residues within the mouse SorCS1b intracellular region (ICR), including T1151 and S1155, in the brain, and identifies GSK3β as the top candidate kinase for T1151 phosphorylation (100% site percentile). Given that SorCS1b OE reduces tau hyperphosphorylation in AD mice, this suggests that the phosphorylated SorCS1b ICR may interact with GSK3β to suppress its kinase activity. Such a mechanism would be analogous to the phosphorylated ICR of LRP6, a Wnt co-receptor, which acts as a “pseudo-substrate” that accesses the catalytic pocket of GSK3β to directly inhibit its activity (*55–58*). Future studies using biochemical experiments are needed to determine how SorCS1 ICR is phosphorylated under physiological and AD conditions, and whether and by what mechanisms the SorCS1 ICR modulates the kinase activity of GSK3β.

Our study highlights several limitations of the 5xFAD model. First, 5xFAD mice in our cohorts exhibited impaired swimming performance, evidenced by reduced velocity, suggesting that the Morris water maze may not have been the most appropriate assay to assess the spatial learning and long-term reference memory of our cohorts. Future studies incorporating less motor-demanding paradigms, such as the Barnes maze or radial arm maze, will be important to determine whether the cognitive benefits of SorCS1 OE extend to measures of spatial learning and reference memory that are less influenced by locomotor performance. Second, the 5xFAD model displays an early onset and rapid, aggressive progression of pathology (*59, 60*), indicating that it would be suitable for modeling early-onset AD but not sporadic, late-onset Alzheimer’s disease (LOAD), which is the most common form (*61*). In addition, transgenic models including 5xFAD have intrinsic limitations related to APP OE, including axonal transport deficits and the overproduction of additional APP fragments that can generate artifact phenotypes (*62*). These issues could be why SorCS1 OE is unable to produce the expected rescue effects, particularly in the Morris water maze test, as we observed. Given these limitations of the 5xFAD line, it would be appropriate to extend this work using a LOAD-relevant knock-in model, such as one expressing human wild-type *APP* under the endogenous mouse *App* promoter (*63*). Third, our study examined SorCS1 OE initiated prior to the onset of histopathological, synaptic, and behavioral phenotypes (*39, 42, 64*). Determining whether SorCS1 OE can also confer benefit after disease onset will be important for establishing its translational relevance and therapeutic potential.

Our findings further identify several important directions for future research. Although SorCS1 OE confers robust protection, the mechanisms underlying its effects on working memory and synapse preservation remain incompletely understood. This likely reflects its multifaceted actions within the crosstalk between amyloid and tau pathologies (*3, 65*): inhibition of β-Nrxn-AβO interaction (*26*), reduction of tau hyperphosphorylation, and promotion of AMPAR trafficking (*27*). Defining how these pathways converge to promote resilience may reveal fundamental mechanisms of synaptic protection and uncover new therapeutic targets downstream of SorCS1. In addition, our gain-of-function studies provide a framework for investigating the physiological role of endogenous SorCS1. Future loss-of-function studies, such as neuronal SorCS1 deletion in AD models, will be essential to determine whether SorCS1 functions as an intrinsic regulator of cognitive resilience and disease progression, and whether enhancing SorCS1 signaling represents a viable disease-modifying strategy. Finally, we serendipitously observed that neuronal SorCS1 OE may reduce microglial activation, suggesting a potential role in coordinating neuronal and immune responses during neurodegeneration. Future studies will be important to establish whether this effect is secondary to preserved synaptic integrity or results from direct modulation of neuron-microglia signaling pathways. Clarifying these mechanisms could extend the significance of SorCS1 beyond synaptic protection and position it as a regulator of neuroimmune homeostasis. Together, our findings provide a framework for understanding how SorCS1 promotes resilience across multiple dimensions of AD pathology and support its potential as a therapeutic target for preserving brain function in neurodegenerative disease.

## Materials and Methods

### Animal and ethics statement

All animal experiments were carried out in accordance with Canadian Council on Animal Care (CCAC) guidelines and were approved by the Institut de Recherches Cliniques de Montréal (IRCM) Animal Care Committee (Protocol number: 2020-10 and 2024-06). Mice were group-housed (two to five per cage) on a 12-h light/dark cycle with ad libitum access to food and water. Sex was considered in the study design. All experiments and data analyses were performed with the experimenters blinded to genotype and treatment.

### Mouse generation and treatment

The inducible SorCS1 transgene knock-in (KI) mouse line (hereafter referred to as SorCS1KI), overexpressing mouse Sorcs1b (NM_021377.3) together with tdTomato, was generated in collaboration with Cyagen Biosciences. Briefly, a donor vector containing the cassette “CAG promoter-loxP-STOP-loxP-Kozak-Sorcs1-IRES-tdTomato-WPRE-rBGpA” flanked by the homology arms of the mouse ROSA26 gene was constructed. This targeting vector was co-injected with CAS9 and a guide RNA (gRNA) targeting the ROSA26 locus (gRNA sequence: CTCCAGTCTTTCTAGAAGAT<u>GGG</u>) into fertilized C57BL/6J mouse eggs to generate KI offspring. F0 founder animals were identified by long-range PCR followed by sequence analysis. The F0 founders were bred to C57BL/6J wild-type (WT) mice to test germline transmission and generate F1 offspring. Positive F1 animals were subsequently backcrossed with C57BL/6J mice for at least four generations.

As an Alzheimer’s disease model mouse line overproducing Aβ, 5xFAD mice were purchased from MMRRC (Strain #: 034840-JAX) (*39*). Because their C57BL/6J x SJL/J hybrid genetic background may carry the retinal degeneration Pde6b^rd1^ mutation (*66*), 5xFAD mice were crossed with C57BL/6J mice for three generations, followed by genotyping to confirm the absence of Pde6b^rd1^ mutation. For forebrain-specific Cre-mediated recombination upon tamoxifen treatment to enable time-dependent expression of the SorCS1 transgene, Camk2a-CreERT2 mice (Strain #: 012362) (*40*) were purchased from Jackson Laboratory. SorCS1KI mice were crossed with 5xFAD mice and subsequently with Camk2a-CreERT2 mice to obtain male and female offspring carrying the following genotypes: heterozygous 5xFAD (5xFAD^WT/Tg^), SorCS1 homozygous SorCS1KI (SorCS1^KI/KI^) and CreER-positive (CreER^+^). These mice were further bred to generate: non-AD mice: 5xFAD^WT/WT^; SorCS1^KI/KI^; CreER^+^ and AD mice: 5xFAD^Tg/Tg^; SorCS1^KI/KI^; CreER^+^. Tamoxifen was administrated by oral gavage (90 μg per g of body weight) in canola oil, or canola oil alone as a vehicle control, for five consecutive days at 1.5 months old.

### Genotyping

Genotyping was performed by PCR using specific primer sets for each allele. For SorCS1, the primers were: WT forward, 5′-CACTTGCTCTCCCAAAGTCGCTC-3′; WT reverse, 5′-ATACTCCGAGGCGGATCACAA-3′; and KI mutant reverse, 5′-GCATCTGACTTCTGGCTAATAAAG-3′. The expected PCR product sizes were 453 bp for the WT allele and 619 bp for the mutant allele. For 5xFAD, the primers were: common forward, 5′-ACCCCCATGTCAGAGTTCCT-3′; WT reverse, 5′-TATACAACCTTGGGGGATGG-3′; and transgene reverse, 5′-CGGGCCTCTTCGCTATTAC-3′. The expected PCR products were 216 bp for the WT allele and 129 bp for the transgene. For detection of the Cre recombinase transgene, the following primers were used: transgene forward, 5′-GACCTGGATGCTGACGAAG-3′; transgene reverse, 5′-AGGCAAATTTTGGTGTACGG-3′; internal positive control forward, 5′-AGTGGCCTCTTCCAGAAATG-3′; and internal positive control reverse, 5′-TGCGACTGTGTCTGATTTCC-3′. The expected PCR product sizes were 200 bp for the Cre transgene and 521 bp for the internal positive control. For the Pde6b^rd1^ mutation, the primers were: WT forward, 5′-TGACAATTACTCCTTTTCCCTCAGTCTG-3′; WT reverse, 5′-TACCCACCCTTCCTAATTTTTCTCACG-3′; and mutant reverse, 5′-GTAAACAGCAAGAGGCTTTATTGGGAAC-3′. The expected PCR product sizes were 410 bp for the WT allele and 561 bp for the *rd1* mutant allele.

### Behavioral tests

In all experiments, male and female mice were analyzed separately due to the higher prevalence of Alzheimer’s disease in women and the more severe Aβ pathology observed in female 5xFAD mice (*50*). Beginning 7 days prior to the first behavioral test, mice in their home cages were transferred daily from the housing room to the experimental room and left for 1 hour to habituate to the novel environment and experimenter handling. During each session, mice were handled by the experimenter for 5 min.

All behavioral experiments were conducted on age- and sex-matched 6 to 6.5-month-old mice during a similar daytime period. The following groups were included: non-AD without SorCS1 overexpression (OE) (Group 1), non-AD with SorCS1 OE (Group 2), AD without SorCS1 OE (Group 3) and AD with SorCS1 OE (Group 4). Before each experiment, mice were transferred to the test room and habituated for at least 1 hour, with the room lighting condition set as specified for each test. All tests were recorded by a color camera with a GigE interface (Basler) and analyzed using EthoVision XT10 (Noldus).

### Y-maze spontaneous alternation test

A Y-shaped maze with three opaque arms 120 degrees apart (LxWxH: 35 × 5 × 20 cm) was used under overhead lighting at 150 lux to assess exploratory behavior. Mice were placed inside one arm facing away from the center and allowed to move through the apparatus for 8 min. Arm entries were scored when all four paws entered the arm.

### Open field test

Under overhead lighting at 150 lux, mice were placed into the center of an open square field (LxWxH: 50 × 50 × 38 cm) at the start of the trial and exploration was recorded for 10 min. To assess locomotor activity, total traveled distance and mean velocity were measured. To assess anxiety-like behavior, time spent in the center area (40% of the total surface) was measured.

### Morris water maze

The apparatus consisted of a circular pool (120 cm in diameter, 50 cm deep) filled with opaque water maintained at 22 ± 1 °C. A hidden platform (10 cm in diameter) was submerged 1 cm below the water surface and placed in the center of the target quadrant. During training sessions, mice underwent four trials per day for five consecutive days, with randomized start positions and an inter-trial interval of at least 10 minutes. Each trial lasted up to 60 seconds, and mice that failed to locate the platform were guided to it and allowed to remain there for 15 seconds. For the probe trial on day 5, a 60-second trial was performed with the platform removed to evaluate memory retention. Time spent in each quadrant and the number of platform crossings were recorded. A visible platform test was conducted following the probe trial to check for visual and motor deficits.

### Crude synaptosome preparation

The crude synaptosome fraction from hippocampi and cortices of 6.5 to 7-month-old mice was purified using a modified protocol as previously described(*67*). Briefly, mice in each group were anesthetized with a ketamine/xylazine mixture (100 mg/kg ketamine, 10 mg/kg xylazine in 0.9% saline) and then decapitated. Immediately afterward, the cortex and hippocampi from each mouse were dissected and separately homogenized in cold buffer A (5 mM HEPES, pH 7.4, 1 mM MgCl_2_, 0.5 mM CaCl_2_, 1 mM DTT, 0.32 M sucrose, supplemented with protease inhibitors (Roche, #5892953001)). The homogenate was centrifuged at 1400 × *g* for 10 min at 4 °C. The resulting pellet was resuspended in buffer A, homogenized again, and centrifuged at 900× *g* for 10 min at 4 °C. The supernatant was combined with the initial supernatant to yield the total lysate fraction. The combined supernatants were centrifuged at 12,000 × *g* for 10 min at 4 °C, and the resulting pellets were resuspended with cold buffer B (6 mM Tris, pH 8.1, 0.32 M sucrose, 1 mM EDTA, 1 mM EGTA, 1 mM DTT, supplemented with protease inhibitors), producing the crude synaptosome preparation. Protein concentrations were measured using DC protein assays (Bio-Rad).

### Immunoblot assay

Protein samples were separated on 8–12% SDS-polyacrylamide gels and transferred onto 0.2 μm polyvinylidene difluoride (PVDF) membranes (Bio-Rad, #1620177). Membranes were blocked with either 5% bovine serum albumin (BSA) in Tris-buffered saline containing 0.1% Tween-20 (TBST) or 5% skim milk in TBST, depending on the antibody, and incubated overnight at 4°C with the following primary antibodies: anti-SorCS1 (1:1,000; rabbit; Abcam, #ab93331), anti-PSD-95 (1:2,000, mouse IgG2a, clone 6G6-1C9, ThermoFisher Scientific, #MA1-045), anti-Synaptophysin (1:1,000, chicken IgY, Synaptic Systems, #101009), anti-pan-Neurexin (1:2,500, rabbit polyclonal, MilliporeSigma, #ABN161-I), anti-Aβ1–16 (1:5,000; mouse IgG1, clone 6E10, BioLegend, #SIG-39300), anti-GluA1 (1:1,000, rabbit polyclonal, MilliporeSigma, #3755420), anti-GluA2 (1:1,000, mouse IgG2a, clone 6C4, Abcam, #MAB397), anti-total Tau (1:2,000, mouse IgG2a, clone PC1C6, MilliporeSigma, #MAB3420), anti-phospho-Tau (Ser202/Thr205) (1:2,000, mouse IgG1, clone AT8, ThermoFisher Scientific, #MN1020), anti-GSK3β (1:1,000, mouse; MilliporeSigma, #05-184-I), anti-phospho-GSK3β (Ser21/9) (1:1,000, rabbit monoclonal; clone D17D2, Cell Signaling Technology, #8566S), and anti-β-actin (1:5,000; mouse; clone 15G5A11/E2; ThermoFisher Scientific, #MA1-140).

Following primary antibody incubation, membranes were washed with TBST and incubated with the appropriate horseradish peroxidase (HRP)-conjugated secondary antibodies (1:5,000, Jackson ImmunoResearch), including donkey anti-mouse IgG (#715-035-151), donkey anti-rabbit IgG (#711-035-152), or donkey anti-chicken IgY, (#703-035-155). Immunoreactive bands were detected using Clarity™ Western ECL Substrate or Clarity Max™ Western ECL Substrate (Bio-Rad) and visualized with a ChemiDoc™ XRS+ Imaging System (Bio-Rad). Band intensities were quantified using ImageJ after background subtraction and normalized to β-actin.

### Immunohistochemistry

Mice (6.5-7 months old) were deeply anesthetized with a ketamine/xylazine mixture (100 mg/kg ketamine, 10 mg/kg xylazine in 0.9% saline) and transcardially perfused with phosphate-buffered saline (PBS), followed by 4% paraformaldehyde (PFA) in PBS. Brains were dissected, post-fixed overnight in 4% PFA at 4°C, washed in PBS, and cryoprotected in 30% sucrose in PBS for 2–3 days at 4°C with gentle agitation. Brains were embedded in Optimal Cutting Temperature (O.C.T.) compound (Tissue-Tek), frozen, and stored at −80°C until sectioning. Coronal brain sections (18 μm) were prepared using a CryoStar™ NX70 cryostat (ThermoFisher Scientific) and mounted onto Superfrost Plus microscope slides (VWR).

Sections were washed three times with PBS and blocked for 1 h at room temperature in blocking solution containing PBS supplemented with 5% bovine serum albumin (BSA), 2.5% normal donkey serum, and 0.25% Triton X-100. For synaptic marker staining, sections were incubated overnight at 4°C with anti-Synaptophysin (1:1,000, chicken IgY, Synaptic Systems, #101009) and anti-PSD-95 (1:1,000, mouse IgG2a, clone 6G6-1C9, ThermoFisher Scientific, #MA1-045). For amyloid plaque staining, sections were incubated with anti-Aβ1–16 (1:3,000, mouse IgG1, clone 6E10, BioLegend, #SIG-39300).

Following primary antibody incubation, sections were washed three times with PBS and incubated for 1 h at room temperature with DAPI (100 ng/mL) together with the appropriate highly cross-adsorbed Alexa Fluor-conjugated secondary antibodies, including donkey anti-mouse Alexa Fluor 488 (1:500, Jackson ImmunoResearch Laboratories, #715-545-150) and donkey anti-chicken Alexa Fluor 680 (1:500; Jackson ImmunoResearch Laboratories; #703-605-155). After three additional PBS washes, sections were mounted with Elvanol mounting medium containing Tris-HCl, glycerol, polyvinyl alcohol, and 2% 1,4-diazabicyclo[2.2.2]octane (DABCO).

### Quantitative fluorescent imaging and image analysis

All imaging and image analysis were done while blind to the experimental conditions. For immunohistochemistry, fluorescent images were acquired on a Leica SP8 confocal microscope with a 10 × 0.30 NA dry objective for quantification analysis of β-amyloid plaque in cortex and hippocampus or 63 × 1.40 NA oil objective for quantification of Synaptophysin and PSD-95 puncta density in the hippocampal CA1 stratum radiatum. All the data for imaging were collected in random order. Analysis was performed using Metamorph 7.8 (Molecular Devices). Quantification of Synaptophysin and PSD-95 puncta density was performed in the same region of interest (ROI). To identify their puncta, images were thresholded by a constant grayscale value equal to the average of the automatically calculated threshold level of all analyzed images. Thresholded puncta were quantified automatically using the integrated morphometric analysis module of MetaMorph. For β-amyloid plaque analysis, cortical and hippocampal ROIs were manually defined using anatomical landmarks. Images were thresholded using a constant grayscale threshold applied uniformly to all sections within the same experiment to identify 6E10-positive plaques. Following thresholding, binary masks were generated, and individual plaques were automatically segmented based on contiguous immunoreactive objects. Touching plaques were separated using the software’s object-separation algorithm when necessary, and obvious artifacts were excluded manually. The total number of plaques within each ROI was quantified and normalized to the ROI area (plaques/μm²). For all quantitative analyses, values were normalized to the mean of the WT control group.

### Electrophysiology

Mice at 6.5 to 7 months old were deeply anesthetized via intraperitoneal injection of ketamine/xylazine mixture (100 mg/kg ketamine, 10 mg/kg xylazine in 0.9% saline) and subsequently decapitated. Brains were rapidly extracted and immersed in ice-cold slicing solution containing (in mM): 2.5 KCl, 0.2 CaCl₂, 4 MgCl₂, 1.25 NaH₂PO₄, 26 NaHCO₃, 10 D-glucose, and 25.2 sucrose, equilibrated to pH 7.35 with 95% O₂/5% CO₂. Acute 350 µm coronal slices encompassing the dorsal hippocampus were prepared using a VT1000S vibratome (Leica Biosystems) and incubated in carbogenated artificial cerebrospinal fluid (ACSF; in mM: 125 NaCl, 2.5 KCl, 2 CaCl₂, 1 MgCl₂, 1.25 NaH₂PO₄, 26 NaHCO₃, 25 D-glucose; pH 7.35). Slices recovered at 32 °C for 1 h, followed by room-temperature (RT) incubation prior to recordings.

For electrophysiology, slices were transferred to a recording chamber and continuously perfused with carbogenated ACSF at RT. Extracellular field excitatory postsynaptic potentials (fEPSPs) were elicited in the CA1 stratum radiatum by Schaffer collateral stimulation using a concentric bipolar microelectrode (FHC Inc.) and recorded with ACSF-filled glass pipettes. Signals were amplified (MultiClamp 700B, Molecular Devices), digitized (Digidata-1550B, Molecular Devices), and acquired with Clampex 11.2; analyses were performed in Clampfit 11.2.

Input–output (I/O) relationships were assessed by plotting fEPSP slopes against fiber volley (FV) amplitudes. Paired-pulse ratios (PPRs) were determined at interpulse intervals of 25, 50, 100, 200, and 500 ms, calculated as the slope of the second fEPSP divided by the first. Statistical analyses included linear regression slope comparison for fEPSP vs. FV amplitude relationships and two-way repeated-measures ANOVA for PPR curves.

### Plasmids

The plasmids pCAG-SorCS1-myc and HA-CD4 were described in our previous studies (*26*). The following plasmids were obtained as kind gifts: HA-tagged GSK3β constructs, including wild-type (WT; Addgene, plasmid #14753), kinase-dead K85A (Addgene, plasmid #14755), and GFP-HA (Addgene, plasmid #229501).

### Co-Immunoprecipitation

HEK293T cells were transfected with the following plasmid combinations: (1) SorCS1-myc alone (a negative control), (2) SorCS1-myc with GFP-HA (an additional negative control), (3) SorCS1-myc with HA-GSK3β wild-type (WT) and (4) SorCS1-myc with HA-GSK3β K85A (kinase-dead mutant). For each condition, 12 μg total DNA was transfected per well using 30 μL of TransIT®-LT1 Transfection Reagent (Mirus Bio. LLC, # MIR2305) and maintained in DMEM supplemented with 10% fetal bovine serum (FBS) for 24 h. The transfected cells were harvested on ice using 1 mL of cold PBS and lysed with vortexing every 5 min in 1300 μl of IP lysis buffer (10 mM HEPES pH 7.4, 150 mM NaCl, 2 mM CaCl_2_, 1 mM MgCl_2_ and 0.1% Tween-20) supplemented with protease and phosphatase inhibitors for 1h at RT. Lysates were incubated with 3 μg anti-HA antibody (Abcam; rabbit polyclonal; #ab9110) for 1h at RT with rotation, followed by incubation with 50 μl of Protein-G magnetic bead slurry (Dynabeads, Invitrogen, #10004D) was added, and the samples were incubated for 1h at 4 °C on a rotating platform. Protein complexes pulled down by the Protein-G beads were purified by magnetic separation (DynaMag-2 Magnet, Invitrogen, #12321D), washing the Protein-G beads once with binding buffer followed by four washes with PBS. Samples containing protein complexes attached to the beads were boiled in 2x Laemmli buffer containing β-mercaptoethanol, resolved by SDS-PAGE, and immunoblotted with anti-myc antibody (Cell Signaling Technology, mouse IgG2a, clone 9B11, #2276S) and anti-HA antibody (Aves Labs, chicken polyclonal, #ET-HA100).

### Statistical analysis

Statistical analyses were performed using GraphPad Prism version 10.2.0 (GraphPad Software, Inc.). Sample sizes were not predetermined using statistical methods. Data were assumed to follow a normal distribution. Statistical comparisons were performed using two-way ANOVA followed by Sidak’s multiple comparison tests or two-tailed unpaired Student’s t-tests, as specified in the figure legends. Data are presented as mean ± SEM, and statistical significance was defined as p < 0.05.

## Acknowledgements

This work was supported by the Canadian Institutes of Health Research (CIHR) grants (PJT-159588 and PTJ-191947) and Fonds de la Recherche du Québec– Santé (FRQS) Research Scholars (Junior 2 (29106) and senior (251655)) to H.T., Alzheimer Society Research Program (ASRP) doctoral fellowship to N.Y., and an IRCM Young Research scholarship and an FRQS Master’s Training scholarship (2005634) to M.W.. N.Y. and H.T. conceived the study, designed the experiments, and wrote the manuscript. N.Y. carried out most of the experimental work, including behavioral experiments immunohistochemistry and biochemical assays. A.K.L. contributed to the generation for SorCS1 KI mouse line. F.B.B. contributed to immunohistochemistry experiments. M.W. contributed to some biochemical experiments. M.I. and H.T. contributed to electrophysiology experiments. H.T. supervised the study. All authors reviewed and approved the final manuscript. The authors declare that they have no competing interests. The data that supports the findings of this study are available within the article and its Supplementary Information file, or from the corresponding author upon request.

## Notes

### Competing Interest Statement

The authors have declared no competing interest.

