## Supplementary Information for "SorCS1 promotes synaptic and cognitive resilience despite amyloid pathology in Alzheimer’s disease model mice"

This document contains:

Supplementary Figures 1 to 3

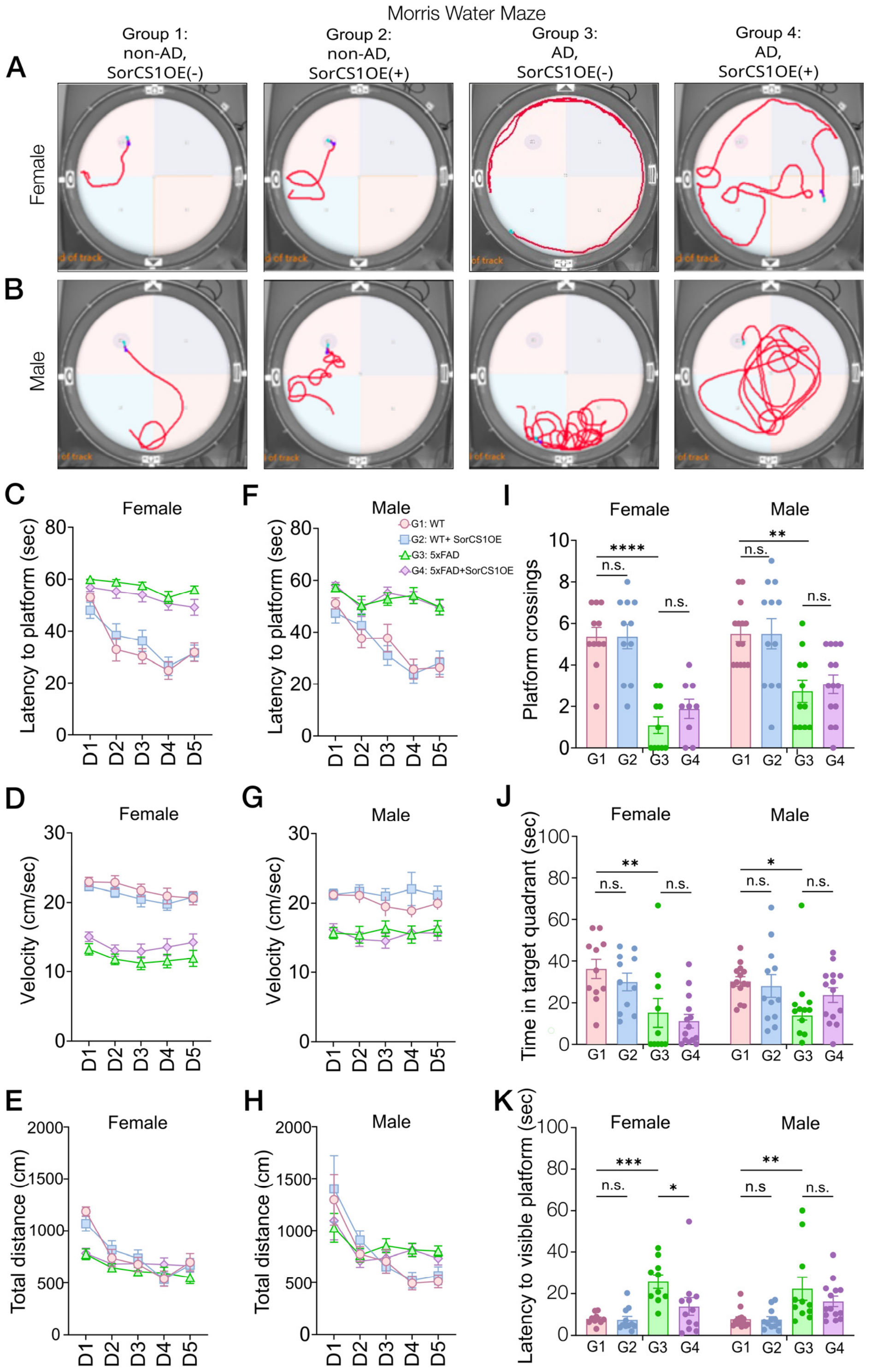


**Supplementary Figure 1. Neuronal SorCS1 overexpression does not improve Morris water maze performance in male or female 5xFAD mice**

**(A, B)** Representative swim trajectories during the probe trial from male **(A)** and female **(B)** mice in Groups 1-4 (Group1: non-AD without SorCS1 overexpression (OE); Group 2: non-AD with SorCS1 OE; Group 3: AD without SorCS1 OE and Group 4: AD with SorCS1 OE).

(**C**-**H**) Spatial learning acquisition during hidden-platform training in male (**C**-**E**) and female (**F**-**H**) mice. Escape latency **(C, F)**, swim velocity **(D, G)**, and total distance traveled **(E, H)** were assessed over five consecutive training days.
(**I**, **J**) Probe trial performance in male and female mice from Groups 1-4. The number of platform crossings **(I)** and time spent in the target quadrant **(J)** were quantified.
(**K**) Performance during cue task (visible platform test) in male and female mice from Groups 1-4.
Data are presented as mean ± SEM. Individual mice are shown as dots where applicable. Acquisition data were analyzed using two-way repeated-measures ANOVA, whereas probe trial and cue task data were analyzed using two-way ANOVA followed by Tukey's multiple comparisons test. Group sizes were as follows: male mice, n = 14 (Group 1), 12 (Group 2), 11 (Group 3), and 13 (Group 4); female mice, n = 15 (Group 1), 15 (Group 2), 12 (Group 3), and 11 (Group 4). *P < 0.05, **P < 0.01, ***P < 0.001, ****P < 0.0001; ns, not significant.

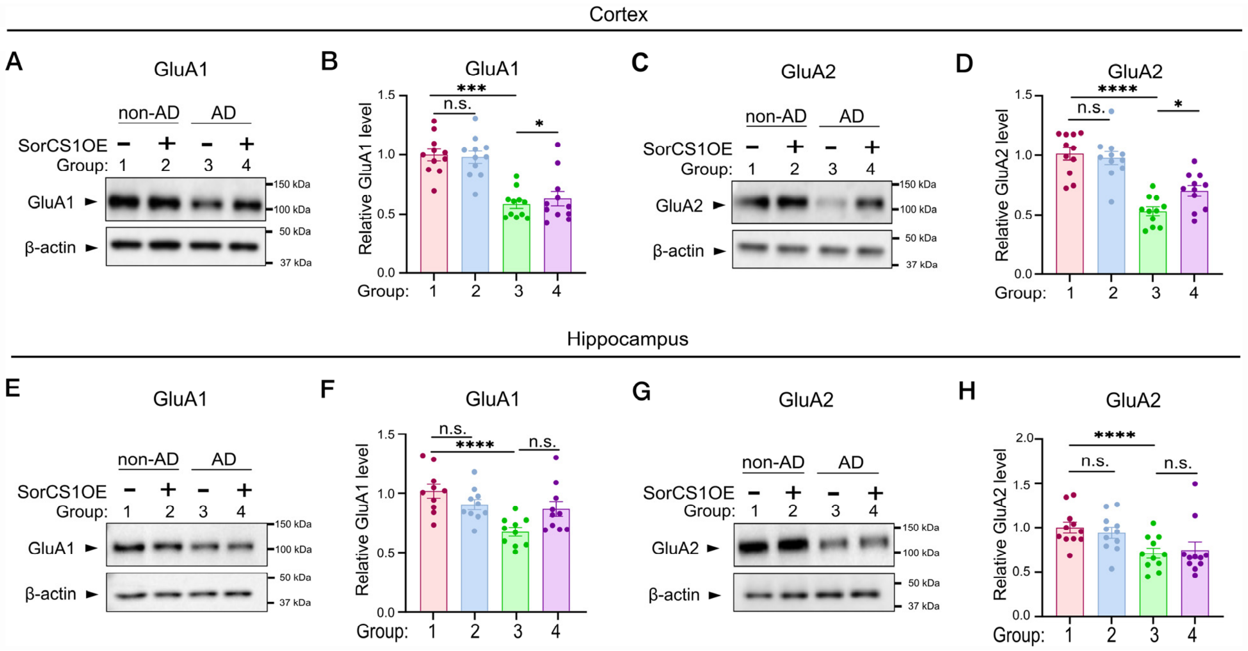


**Supplementary Figure 2. Neuronal SorCS1 overexpression restores synaptosomal levels of AMPA receptor subunits GluA1 and GluA2 in the cortex, but not in the hippocampus**

(**A**, **C**, **E**, **G**) Representative immunoblots of GluA1 (**A**, **E**) and GluA2 (**C**, **G**) in cortical (**A**, **C**) and hippocampal (**E**, **G**) synaptosomal fractions isolated from female mice in Groups 1-4.
(**B**, **D**, **F**, **H**) Quantification of GluA1 (**B**, **D**) and GluA2 (**F**, **H**) protein levels in cortical (**B**, **D**) and hippocampal (**F**, **H**) synaptosomal fractions from Groups 1-4. Protein expression was normalized to β-actin and expressed relative to the mean value of Group 1. Data are presented as mean ± SEM. Statistical analyses were performed using two-way ANOVA followed by Dunnett's multiple comparisons test. Group sizes: n = 10 mice per group for cortical analyses and n = 11 mice per group for hippocampal analyses. *$P < 0.05$, **$P < 0.01$, ***$P < 0.001$, ****$P < 0.0001$; ns, not significant.

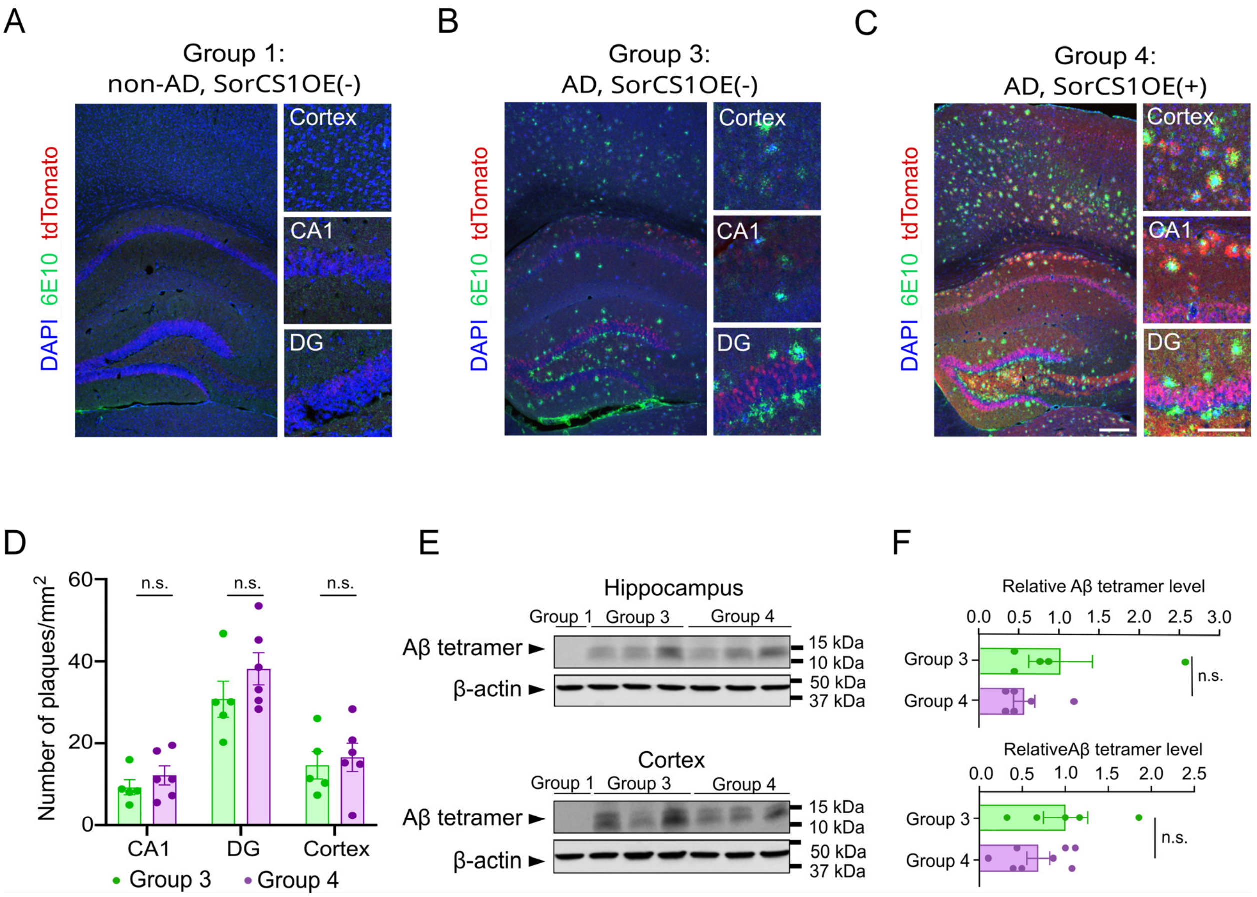


**Supplementary Figure 3. Neuronal SorCS1 overexpression does not affect amyloid plaque burden or Aβ tetramer levels in male 5xFAD mice**

(**A-C**) Representative immunofluorescence images of amyloid plaques labeled with 6E10 antibody (green) in male mice from Groups 1, 3 and 4. tdTomato fluorescence, indicating transgene expression, is shown in red, and nuclei were counterstained with DAPI (blue). Higher-magnification images of the hippocampal CA1 region, and dentate gyrus (DG), and cortex are shown on the right. Scale bars: 500 µm (low magnification) and 250 µm (high magnification).

(**D**) Quantification of 6E10-positive plaque density (plaques/mm²) in the CA1, DG and cortex of male mice from Groups 3 and 4. No significant differences in plaque density were detected between Groups 3 and 4 in any brain region examined. Data are presented as mean ± SEM. Statistical analyses were performed using unpaired two-tailed Student's *t*-tests. Group sizes: n = 5 and 6 mice for Group 3 and Group 4, respectively. ns, not significant.

(**E, F**) Representative immunoblots of (**E**) and quantification (**F**) of Aβ tetramers in hippocampal (top) and cortical (bottom) lysates from Group 3 and 4 male mice. β-actin served as the loading control. Quantification revealed no significant differences in Aβ tetramer abundance between the two groups in either brain region. Data represent mean ± SEM. Statistical analyses were performed using unpaired two-tailed Student's *t*-tests. Group sizes: n = 5 and 6 mice for Group 3 and Group 4, respectively. ns, not significant.
